# The PAR2 signal peptide prevents premature receptor cleavage and activation

**DOI:** 10.1101/760769

**Authors:** Belinda Liu, Grace Lee, Jiejun Wu, Janise Deming, Chester Kuei, Anthony Harrington, Lien Wang, Jennifer Towne, Timothy Lovenberg, Changlu Liu, Siquan Sun

## Abstract

Unlike closely related GPCRs, protease-activated receptors (PAR1, PAR2, PAR3, and PAR4) have a predicted signal peptide at their N-terminus, which is encoded by a separate exon, suggesting that the signal peptides of PARs may serve an important and unique function, specific for PARs. In this report, we show that the PAR2 signal peptide, when fused to the N-terminus of IgG-Fc, effectively induced IgG-Fc secretion into culture medium, thus behaving like a classical signal peptide. The presence of PAR2 signal peptide has a strong effect on PAR2 cell surface expression, as deletion of the signal peptide (PAR2ΔSP) led to dramatic reduction of the cell surface expression and decreased responses to trypsin or the synthetic peptide ligand (SLIGKV). However, further deletion of the tethered ligand region (SLIGKV) at the N-terminus rescued the cell surface receptor expression and the response to the synthetic peptide ligand, suggesting that the signal peptide of PAR2 may be involved in preventing PAR2 from intracellular protease activation before reaching the cell surface. Supporting this hypothesis, an Arg36Ala mutation on PAR2ΔSP, which disabled the trypsin activation site, increased the receptor cell surface expression and the response to ligand stimulation. Similar effects were observed when PAR2ΔSP expressing cells were treated with protease inhibitors. Our findings indicated that these is a role of the PAR2 signal peptide in preventing the premature activation of PAR2 from intracellular protease cleavage before reaching the cells surface. The same mechanism may also apply to PAR1, PAR3, and PAR4.

## Introduction

G-protein coupled receptors (GPCRs) are a class of 7 transmembrane domain cell surface receptors and consist of the largest receptor family in mammals and other organisms. They are involved in the signal transduction of almost every system in human physiology, including the sensory, metabolic, endocrine, immune, and the nervous systems. Unlike many other cell surface receptors that have a classical signal peptide to lead the proteins to cells surface, the majority of GPCRs (>90%) do not have a signal peptide (1). In general, class B receptors such as the secretin receptor (2), corticotropin-releasing hormone (CRH) receptors (3), the Glucagon receptor (4), and Glucagon-like peptide receptors (5) and the class C GPCRs, such as metabotropic glutamate receptors (6), GABA receptors (7), and adhesion GPCRs (8), which have relatively large N-terminal extracellular domains are more likely to have signal peptides than class A receptors (**Fig 1A**). It is hypothesized that the presence of the signal peptide helps the large hydrophilic N-terminus to cross the plasma membrane. Most class A GPCRs do not have classical signal peptides. It is believed that the first transmembrane domain of these class A GPCRs serves as a signal anchor sequence allowing these receptors to translocate to the cell membrane after translation and assembly in the endoplasmic reticulum (ER) (9). Protease-activated receptors (PARs), including PAR1, PAR2, PAR3, and PAR4 belong to class A GPCR receptor sub-family (10). Homology-wise, they are very closely related to cysteinyl leukotriene receptors (CYSLT), niacin receptors (GPR109), lactic acid receptor (GPR81), and the succinate receptor (GPR91). Unlike their closest neighbors (**Fig 1B**), which do not possess a signal peptide, all PARs have a predicted signal peptide at their N-termini (**Fig 1C**). Genomic analyses show, in contrast to their closest neighbors that are all encoded by single exon genes, PARs have an additional exon encoding only the signal peptides (**Fig 1C**), suggesting that these signal peptides may play a specific role for PARs. In this report, we used PAR2 to study the importance of the signal peptide in PAR receptor function and localization.

**Fig 1.**
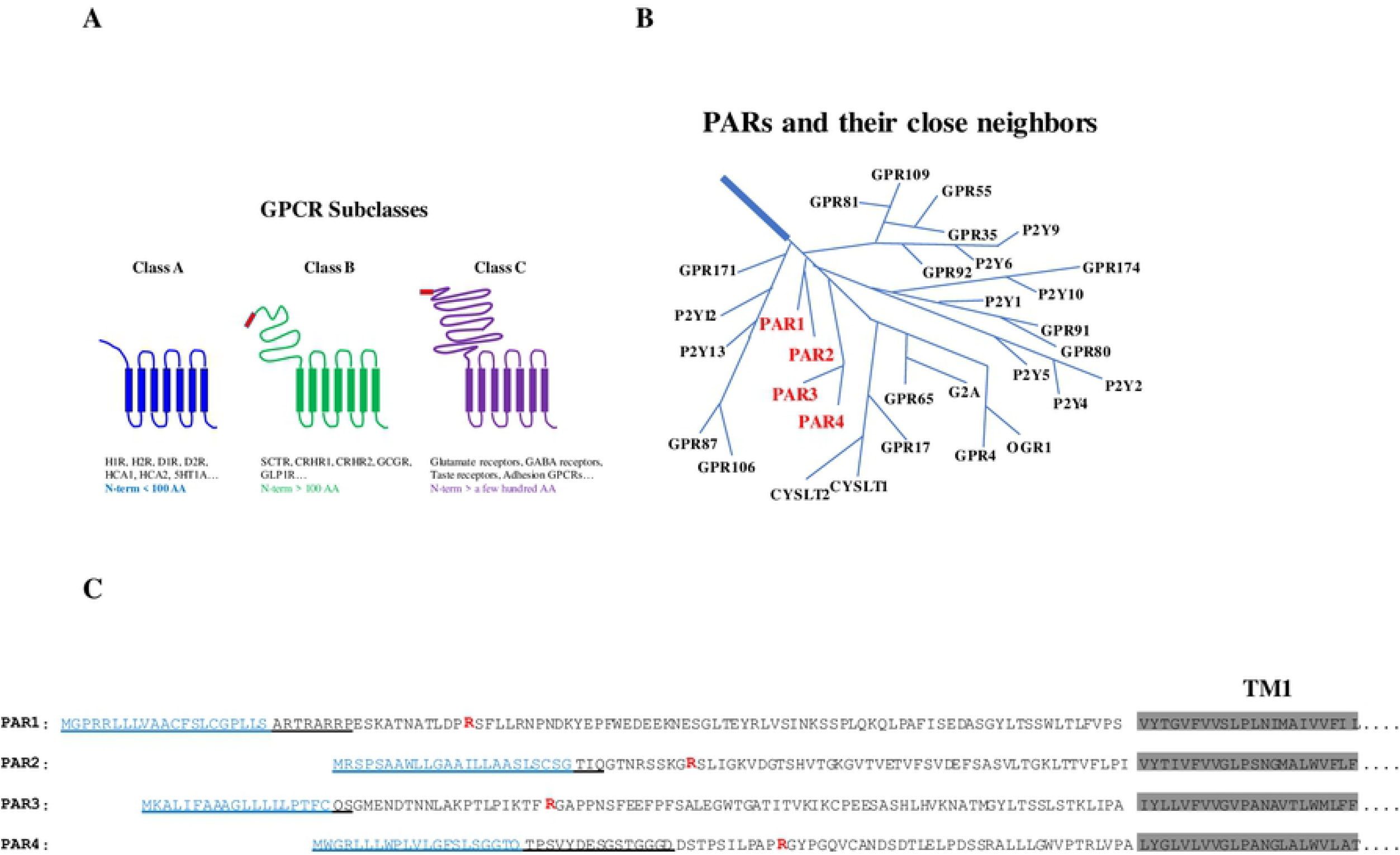
PAR receptors are unique group of receptors in the Class A subfamily. **A**. Examples of GPCR subfamily members and signal peptide possession. The signal peptide regions in Class B and C are shown in red. **B**. PAR receptors and their closest neighbors, grouped by sequence similarity. **C**. The N-terminal amino acid sequence of PAR1-4. The signal peptides are shown in blue. Each PAR receptor is encoded by 2 exons. The protein regions coded by the first exons are underlined. The Arg residues involved in receptor cleavage and activation are shown in red. The first transmembrane domains are shaded.

## Materials and methods

### Reagents

The PAR2 agonist peptide ligand, SLIGKV, was synthesized by Innopep, Inc. (San Diego, CA). Trypsin (sequencing grade), thrombin, and protease inhibitors were purchased from Sigma Aldrich (St. Louis, MO).

#### Quantitative PCR analysis of the mRNA expression levels of PARs

Total RNAs were isolated from COS7, HEK293, and CHO-K1 cells respectively using a RNA isolation kit (RNeasy Mini Kit) from Qiagen. cDNAs were synthesized from the isolated RNA using a cDNA synthesis kit (Advantage RT-PCR kits) from Clontech (Mountain View, CA). Specific primers designed according to human, monkey, and hamster PAR1, PAR2, PAR3, and PAR4 were used to quantify each mRNA expression using a qPCR machine (QuantStudio, ABI) as described (11). In parallel, primers for β-actin were used to amplify β-actin cDNA as the internal controls. The relative expressions of different PAR mRNAs were normalized using the expression level of β-actin. The qPCR primers were designed based on the published cDNA sequences and the primer sequences are listed in **Table 1**.

**Table 1.**
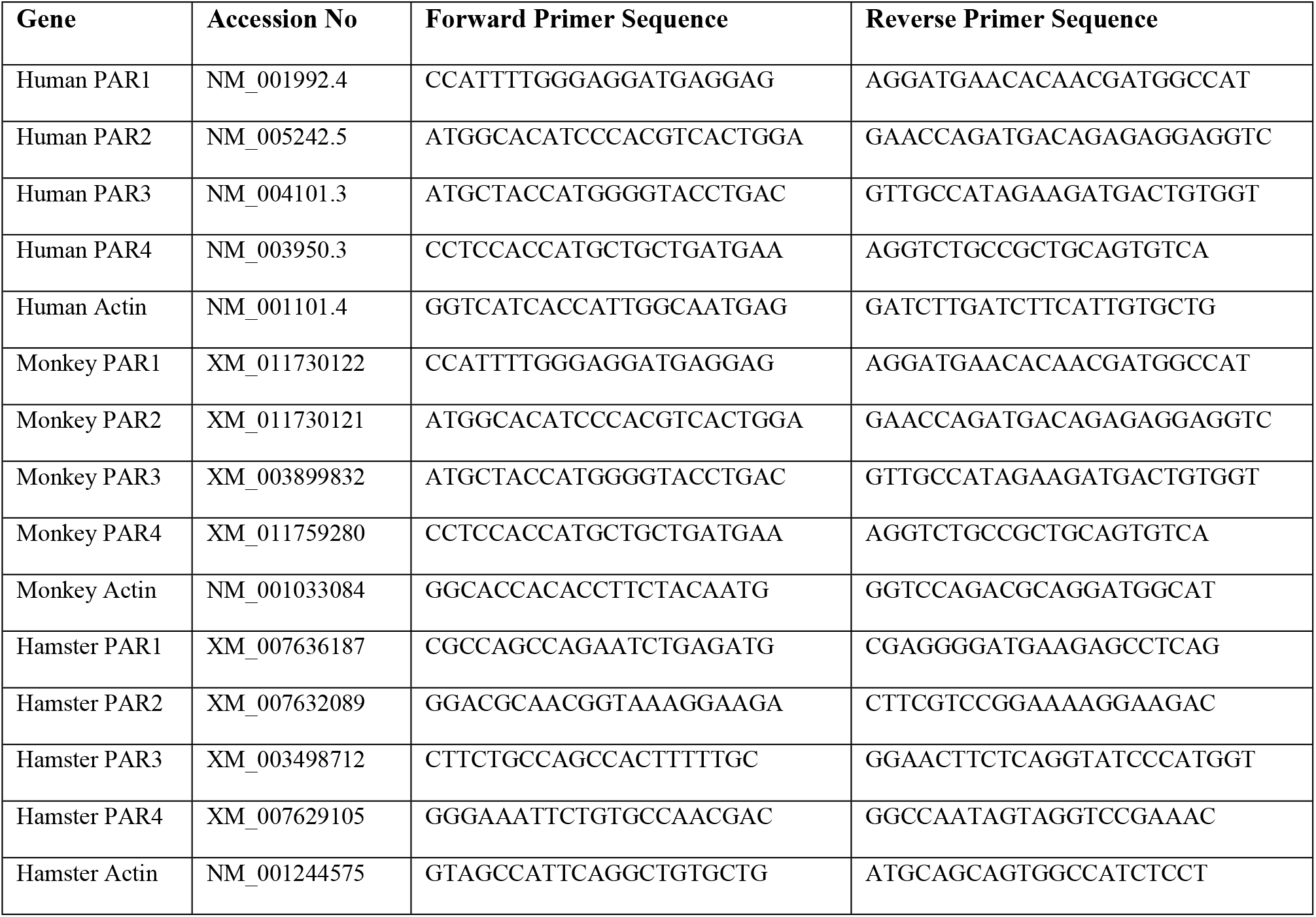
The Genbank accession numbers for cDNA sequences used for PCR primer design and primer sequences for qPCRs. All primer sequences are shown from 5’ to 3’.

#### Generation of PAR1, PAR2 knock-out cell line

A PAR1, PAR2 knock-out HEK293 cell line was created by Applied StemCells (Milpitas, CA) using a CRISPR/Cas9 approach. Briefly, the coding region (nucleotide 374-643) of PAR1, which encodes the protein region TM2 to TM3 of PAR1, was deleted. Similarly, the coding region (281-627) of PAR2, which encodes the protein region TM2 to TM3 of PAR2, was deleted. Single cell clones were isolated. PCR analysis of the genomic DNA by PCR followed by DNA sequencing were used to confirm the deletion of the DNA fragments.

#### Molecular cloning of PAR2 constructs

The PAR2 coding region was amplified by polymerase chain reaction (PCR) using primers (5’ atg tct GAA TTC GCC ACC atg cgg agc ccc agc gcg gcg tgg ctg ctg −3’; reverse primer: 5’-atg tct GCG GCC GCt caa tag gag gtc tta aca gtg gtt gaa ct-3’) designed based on the published PAR2 coding sequence (Genbank Accession No. NM_005242.5). Human colon cDNA purchased from Clontech (Palo Alto, CA) was used as the template. Expanded high fidelity PCR system (Roche Life Science, Indianapolis, IN) was used to amplify the full length PAR2 cDNA coding region. The resulting DNA was digested using EcoR1 and Not1 restriction enzymes (Promega, Madison, WI) and then cloned into pcDNA3.1 (Invitrogen, Carlsbad, CA). The insert region was then sequenced by Eton Biosciences (San Diego, CA) and the identity of the entire coding region was confirmed.

Expression constructs for PAR2 with an Arg36Ala mutation (PAR2(R36A)), PAR2 without the signal peptide (PAR2ΔSP), PAR2ΔSP with an Arg36Ala mutation (PAR2ΔSP(R36A)), and PAR2 without the signal peptide and the tethered ligand (PAR2ΔSPΔL) were generated by site directed mutagenesis using overlapping PCR approach (12).

Genes for PAR2 with the signal peptide coding regions replaced by the insulin signal peptide, or the insulin receptor signal peptide were synthesized by Eton Biosciences. Similarly, expression constructs for various PAR2 variant with a GFP fused to the C-termini, human IgG-Fc coding region with or without a PAR2 signal peptide coding region, with an insulin, or with an insulin receptor signal peptide coding region were synthesized. The genes were cloned into pcDNA3.1 and the entire coding regions were sequenced to confirm the identities.

#### Intracellular Ca^2+^ mobilization assay

FLIPR-Tetra (Molecular Device, San Jose, CA) was used to monitor intracellular Ca^2+^ mobilization in HEK293 cells, HEK293 cells with PAR1 and PAR2 knocked-out, and cells transiently transfected with various PAR2 expression constructs. Cells were grown in 96-well polyD-lysine coated black FLIPR plates (Corning) in DMEM supplemented with 10% FBS, 1 mM pyruvate, 20 mM HEPES, at 37°C with 5% CO_2_. For transient transfection, cells were grown in 96-well polyD-lysine coated black FLIPR plates and transfected using FuGENE HD (Promega, Madison, WI) as the transfection reagent according to the manufacturer’s instructions. For samples treated with protease inhibitors, protease cocktail was added to cell culture one day after transfection and incubated overnight. Two days after transfection, cell culture media were removed, and cells were washed using HBSS buffer plus 20 mM HEPES. Ca^2+^ dye (Flura 3) diluted in HBSS buffer plus 20 mM HEPES was used to incubate cells at RT for 40 min to allow Ca^2+^ to enter cells. Intracellular Ca^2+^ mobilization stimulated by various concentrations of ligands (trypsin, or peptide ligand) was monitored by FLIPR-Tetra as described (13). The untransfected cells were used as negative controls.

#### Enzyme linked immunosorbent assay (ELISA) for the measurement of IgG-FC secretion

COS7 cells were grown in 6 well plates with DMEM supplemented with 10% FCS, 1 mM pyruvate, 20 mM HEPES, at 37°C with 5% CO_2_ and transfected by different expression constructs for human IgG-Fc with various signal peptide coding regions using LipofectAmine (Invitrogen, Calsbad, CA) as the transfection reagent according to the manufacturer’s instructions. Untransfected cells were used as negative controls.

To measure the secreted human IgG-FC in the medium, one day after transfection, the cells were washed 3 times using PBS and then cultured in serum free DMEM plus 1 mM pyruvate and 20 mM HEPES. Three days after transfection, the conditioned media from the transfected cells were harvested and centrifuged at 10,000 g at 4°C for 20 min to remove the cell debris. 50 ul of the conditioned medium from each transfection was incubated in one well of a 96-well ELISA plate (UltraCruz^®^ ELISA Plate, high binding, 96 well, Flat bottom, Santa Cruz Biotechnology) at 37°C for 1 hr to allow protein in the media to adsorb to the plates. The plates were washed 3 times using PBS + 0.1% Tween-20 (PBST) and blocked using 3% no-fat milk in PBST for 30 min at RT and then incubated using HRP-conjugated goat-anti-human Ig-GF antibody (50 ng/ml) diluted in 3% no-fat milk in PBST at 4°C overnight. The plates were washed 3 times using PBST and then developed using an ELISA developing kit (BD Biosciences). The optical densities at 450 nm were read using an ELISA plate reader (Molecular Devices).

To measure intracellular IgG-Fc protein, one day after transfection, cells were trypsinized and seeded in 96-well culture plates (30,000 cell/well) and grown in DMEM supplemented with 10% FCS, 1 mM pyruvate, 20 mM HEPES. Three days after transfection, the media were removed, and cells were washed using PBS, and then fixed by 10% formaldehyde in PBS at RT for 15 min. The cells were penetrated using 1% Triton-X-100 at RT for 10 min and blocked using 3% no-fat milk in PBST for 30 min at RT. The cells were then incubated using HRP-conjugated goat-anti-human IgG-Fc antibody and the plate was developed and read as described above.

#### Immuno-fluorescent staining of intracellular IgG-Fc

COS7 cells were transfected with various IgG-Fc expression constructs. One day after transfection, cells were trypsinized and seeded in a 4-well cell culture chamber slides (Stellar Scientific, Baltimore, MD) (60,000 cells/well). Three days after transfection, the media were removed, and cells were washed using PBS, and then fixed by 10% formaldehyde in PBS at RT for 15 min. The cells were penetrated using 1% Triton-X-100 at RT for 10 min and blocked using 3% no-fat milk in PBST for 30 min at RT. The cells were then incubated using FITC-labelled goat-anti-human IgG-FC antibody (ThermoFisher Scientific) (200 ng/ul) diluted in 3% no-fat milk in PBST at 4°C overnight. The slides were then washed 3 times using PBST and then dried using cool air and viewed under a fluorescent microscope.

#### Identification of the signal peptide cleavage site of PAR2

COS7 cells were grown in 15 cm dishes in DMEM supplemented with 10% FCS, 1 mM pyruvate, 20 mM HEPES, at 37°C with 5% CO_2_ and transfected by the expression construct of human IgG-FC with the N-terminus of PAR2 using LipofecAmine. One day after transfection, the cells were washed 3 times using PBS and then cultured in serum free Opti-MEM (Life Technology) plus Pen/Strep. Three days after transfection, the media were collected and centrifuged to remove the cell debris. The supernatants were passed through a Protein A (Sigma) affinity column. The column was washed with PBS and eluted using 0.1 M Glycine/HCl (pH 2.8) and then neutralized using 1 mM Tris-HCl, pH 8.0. The eluted protein was first treated with PNGase-F (Promega) to remove the N-linked glycosylation and then analyzed by mass spectrometry to determine the N-terminal sequence. Protein sequencing was performed using a generic insolution protein digestion and LC-MS/MS method. Briefly, a 10 ul protein sample in 50 mM ammonia bicarbonate buffer (pH 7.8) was reduced by 11.3 mM dithiothreitol at 60°C for 30 min (without urea), alkylated with 37.4 mM iodoacetamide (RT, 45 min), and then digested with 0.2 ug Trypsin (37°C, overnight). LC/MS analysis was carried out on an Agilent 1290 UHPLC coupled to a 6550 qTOF mass spectrometer, under the control of MassHunter software version 4.0. Chromatography was run with an Agilent AdvanceBio Peptide Map column (2.1 x 100 mm, 2.7 μm) using water/acetonitrile/0.1% formic acid as mobile phases, and mass spectrometric data were acquired in both MS and MSMS modes.

#### Protease inhibitor treatment of cells recombinantly expressing PAR2 receptors

The wild type and various mutant PAR2 variant expression constructs were transiently transfected into HEK293 cells with *par1* & *par2* genes knocked-out. 24 hrs after transfection, cells were treated for 12 hrs with a protease inhibitor cocktail including 4-(2-aminoethyl)benzenesulfonyl fluoride hydrochloride (AEBSF, 500 uM), Leupeptin (50 uM), aprotinin (50 uM).

#### Measurement of the total and cell surface expression of PAR2 by ELISA

ELISA was used to measure the total and cell surface PAR2 protein expression. The wild type and different mutant PAR2 variants were transiently expressed in HEK293 cells with the endogenous PAR1 and PAR2 knocked-out. The cells were transfected in 10 cm cell culture dishes and, 24 hr after transfection, split into 96-well polyD-lysine coated plate. 48 hr post transfection, cells were fixed as described above. To measure the total PAR2 expression, the fixed cells were penetrated using 1% triton-X-100, and blocked with 3% no-fat milk, and then incubated with a monoclonal antibody [3 μg/ml, mouse anti-human PAR2 (BioLegand, San Diego, CA)], which recognizes the N-terminal region (amino acid residues 37-62) of the human PAR2, at 4°C overnight. The plate was washed with cold PBS 3 times and then incubated using a HRP-conjugated goat-anti-mouse IgG secondary antibody (30 ng/ml, Pierce) at RT for 1 hr. The plate was washed again using PBS and developed using an ELISA developing kit as described above. To measure the cell surface PAR2 expression, the ELISA assays were performed in the same manner as the total PAR2 measurement without using triton-X-100 as the cell penetrating agent.

#### Measurement of the total expression and cellular localization of PAR2-GFP fusion proteins

GFP fusion proteins of PAR2 wild type and various mutants were transiently expressed in 96-well poly-D-lysine plates in HEK293 cells with the endogenous PAR1 and PAR2 knocked-out as described above in methods for *Intracellular Ca^2+^ mobilization assay*. 48 hrs after transfection, the media were aspired, and cells were fixed using 4% Paraformaldehyde in PBS (Sigma). The fluorescent intensities of the cells were read using an Envision plate reader (PerkinElmer). The fixed cells were then analyzed using a confocal microscopy for PAR2 cellular localizations.

## Results and discussions

### PAR2 signal peptide behaves as a classical signal peptide

#### PAR2 signal peptide leads IgG-Fc fragment secretion to the medium

A classical signal peptide is typically found at the N-termini of either secreted proteins (such as insulin) or cell surface proteins (such as insulin receptor). It typically consists of a stretch of 20-30 hydrophobic amino acid residues. Its known function is to help a secreted or a cell surface protein to target the ER during protein translation and cross the plasma membrane. PAR2 has a predicted signal peptide sequence at its N-terminus and we hypothesized that it may function as a classical signal peptide. To address this, we devised a few expression constructs (**Fig 2A**) to test whether the signal peptide of PAR2 enables the secretion of human IgG-Fc fragment (which lacks a signal peptide) into the cell culture medium. IgG-Fc was used since it was easy to detect using ELISA or immune-staining. When recombinantly expressed in mammalian cells, without a signal peptide, IgG-Fc is only expressed intracellularly. In contrast, with a signal peptide, IgG-Fc can be secreted into the cells culture medium. One IgG-Fc construct contained the N-terminus of PAR2 with its signal peptide and another with the PAR2 N-terminus in which its signal peptide was deleted. As positive controls, constructs with an insulin signal peptide (a secreted protein signal peptide) or an insulin receptor signal peptide (a cell surface receptor signal peptide) fused to human IgG-Fc were also used in the experiment. We used immuno-staining (**Fig 2B**) and ELISA (**Fig 2C**) to detect and measure IgG-Fc expression in the transfected cells and demonstrated that all cells transfected with various IgG-Fc expression constructs expressed IgG-Fc in the cells. Consistent with our hypothesis, our results showed that fusing the N-terminus of PAR2 with its signal peptide to the human IgG-Fc, effectively led to the secretion of IgG-Fc to the medium, thus functioning similarly to that of the insulin signal peptide or the insulin receptor signal peptide (**Fig 2D**). In contrast, fusing the PAR2 N-terminus without its signal peptide failed to lead to the secretion of IgG-Fc into the medium.

**Fig 2.**
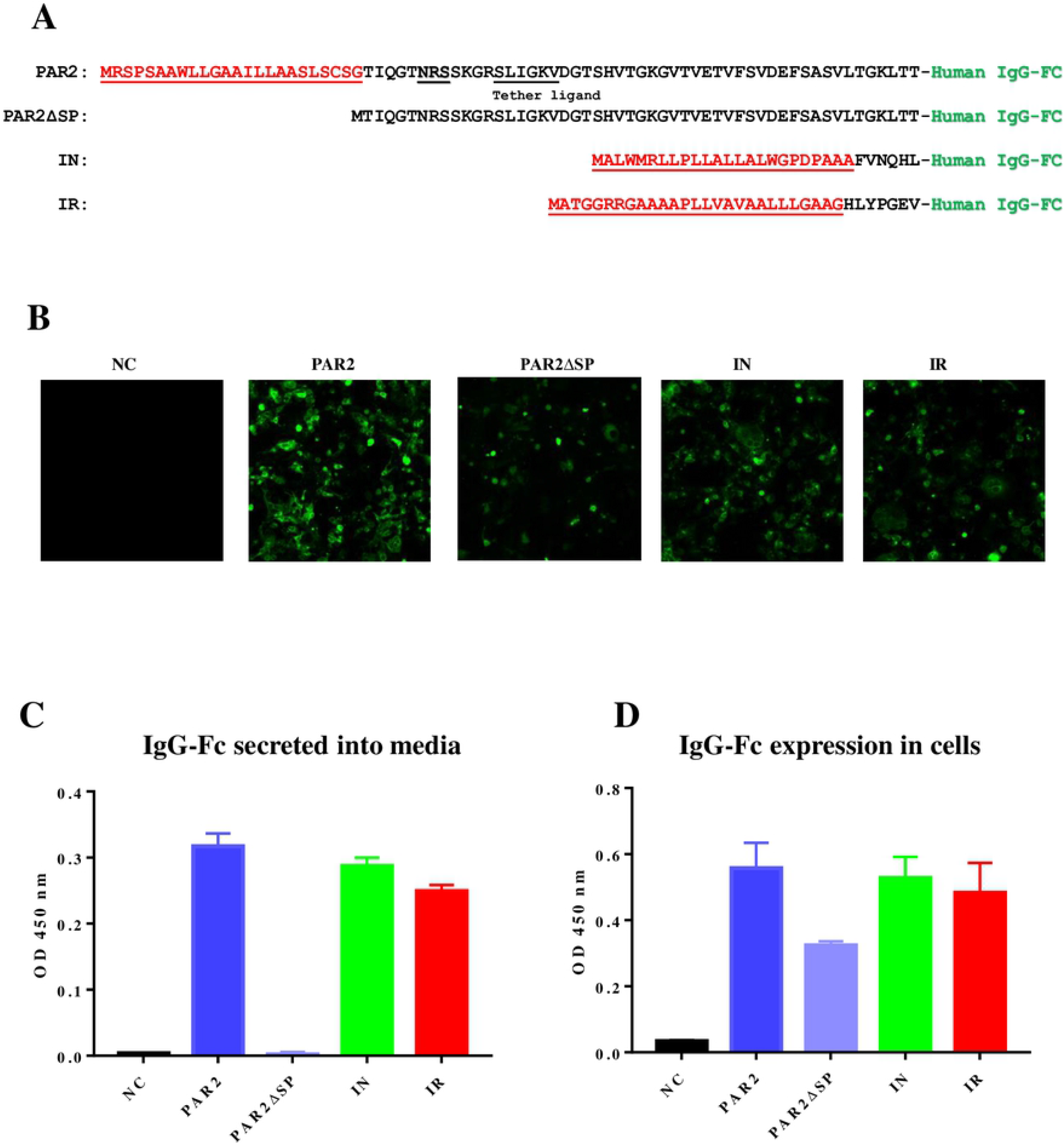
PAR2 signal peptide behaves like a classical signal peptide. A. Expression constructs for testing the roles of the signal peptide of PAR2 in leading IgG-Fc secretion. The N-terminus of PAR2 with its signal peptide (**PAR2**), the N-terminus of PAR2 without the signal peptide (**PAR2ΔSP**), the N-terminus of insulin (**IN**), and the N-terminus of insulin receptor (**IR**) are fused to the human IgG-Fc fragment respectively. The signal peptide regions of PAR2, insulin, and insulin receptor are highlighted in red and underlined. Human IgG-Fc fragment is highlighted in green. B, C. Detection of IgG-Fc expression in cells by immuno-fluorescent staining and ELISA. COS7 cells expressing various IgG-Fc fusion proteins as indicated were fixed, penetrated using detergent, and then detected or stained by FTIC-labeled fluorescent antibodies (**B**) or by ELISA (**C**). D. Detection of IgG-Fc secretion into media by ELISA. Serum free conditioned medium from COS7 cells expressing various IgG-Fc fusion proteins with different N-termini, including PAR2 N-terminus (**PAR2**), PAR2 N-terminus without the signal peptide (**PAR2ΔSP**), the N-terminus of insulin (**IN**), and the N-terminus of insulin receptor (**IR**). Untransfected cells were used as the negative control (**NC**). The experiments have been performed 3 times and very similar results have been observed. Example data is shown.

#### PAR2 signal peptide is cleaved from the mature protein

It has been reported that for the CRF2(a) receptor, the signal peptide may not be cleaved from the mature proteins following membrane insertion (14). To determine if this is the case for the PAR2 signal peptide, we examined whether the signal peptide of PAR2 is cleaved from the mature IgG-Fc protein with the PAR2 N-terminus following secretion. The conditioned medium from the COS7 cells transfected with the expression construct for PAR2 N-terminus fused to IgG-Fc (**Fig 2A**) was collected. Secreted PAR2-IgG-Fc fusion protein was affinity purified, glycosylation moieties removed, and then analyzed by mass spectrometric (MS) protein sequencing. Our results showed that the most N-terminal sequence that matches PAR2 sequence is TIQGTNR (**Fig 3**), suggesting that the signal peptide has been cleaved following the protein secretion, with the cleavage site being between residues Gly^24^ and Thr^25^. Interestingly, a variant sequence TIQGT<u>D</u>R was also observed. This sequence differs from TIQGTNR by one residue (from N to D). Since the residue Asn^30^ is a part of a NRS sequence (a N-linked glycosylation site, **Fig 3**) and glycosylated Asn residues are converted to Asp after de-glycosylation by PNGase-F, our results suggest that at least part of the expressed protein is glycosylated at this N-linked glycosylation site.

**Fig 3.**
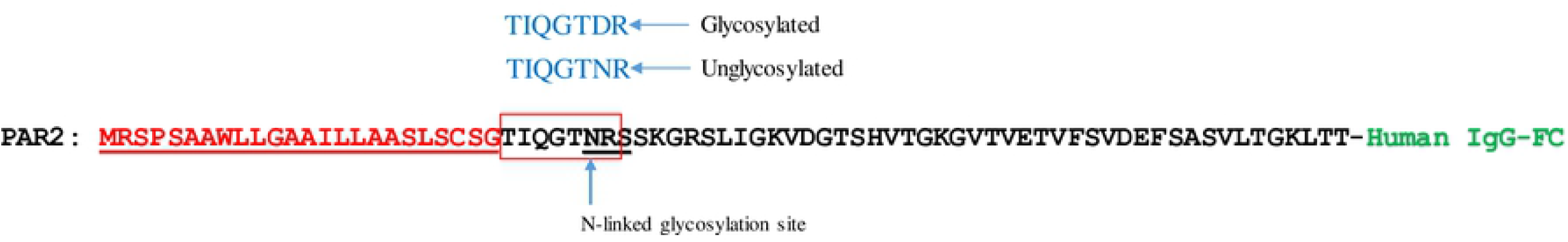
Determination of the N-terminal sequence of PAR2 mature protein. The N-terminal extracellular region of PAR2 is fused to the N-terminus of IgG-Fc. The predicted signal peptide of PAR2 is shown in red and underlined. The IgG-Fc region is shown in green. The potential N-linked glycosylation site, NRS, is underlined. The protein is expressed in COS7 cells and affinity purified. The N-terminus of the purified protein was determined by MS sequencing after trypsin digestion. Two sequences have been observed: TIQTNR and TIQTDR representing unglycosylated and glycosylated PAR2 N-termini.

### PAR2 signal peptide is important for PAR2 receptor functional expression and activation by its ligands

#### Generation of a PAR1 & PAR2 knock-out HEK293 cell line for recombinant expression and characterization of the PAR2 receptor

To evaluate the receptor localization and function of recombinant PAR2, it is essential to have a host mammalian cell line that does not express endogenous PAR2 or other PAR receptors. We tested a few commonly used mammalian cells for recombinant expression, including HEK293, CHO-K1, and COS7 cells and found that all three cell lines express relatively high PAR1 and PAR2 mRNA (**Fig 4A**). In addition, in functional assays, they all responded to PAR1 and PAR2 ligands (thrombin and trypsin, respectively) (**Fig 4B-D**). Since the presence of naturally expressed PAR1 and PAR2 in these host cells would complicate the characterization of the recombinantly expressed PAR2, we created a HEK293 cell line (which does not express PAR3 and PAR4) with both *par1* and *par2* genes knocked-out by CRISPR/cas9 (**Fig 4E**). Pharmacological characterization of this cells line demonstrated that the loss of both *par1* & *par2* led to a lack of responses to PAR1 ligand (thrombin) or PAR2 ligand (trypsin) stimulation (**Fig 4F**). These cells were then used to study expression and localization of recombinant PAR2.

**Fig 4.**
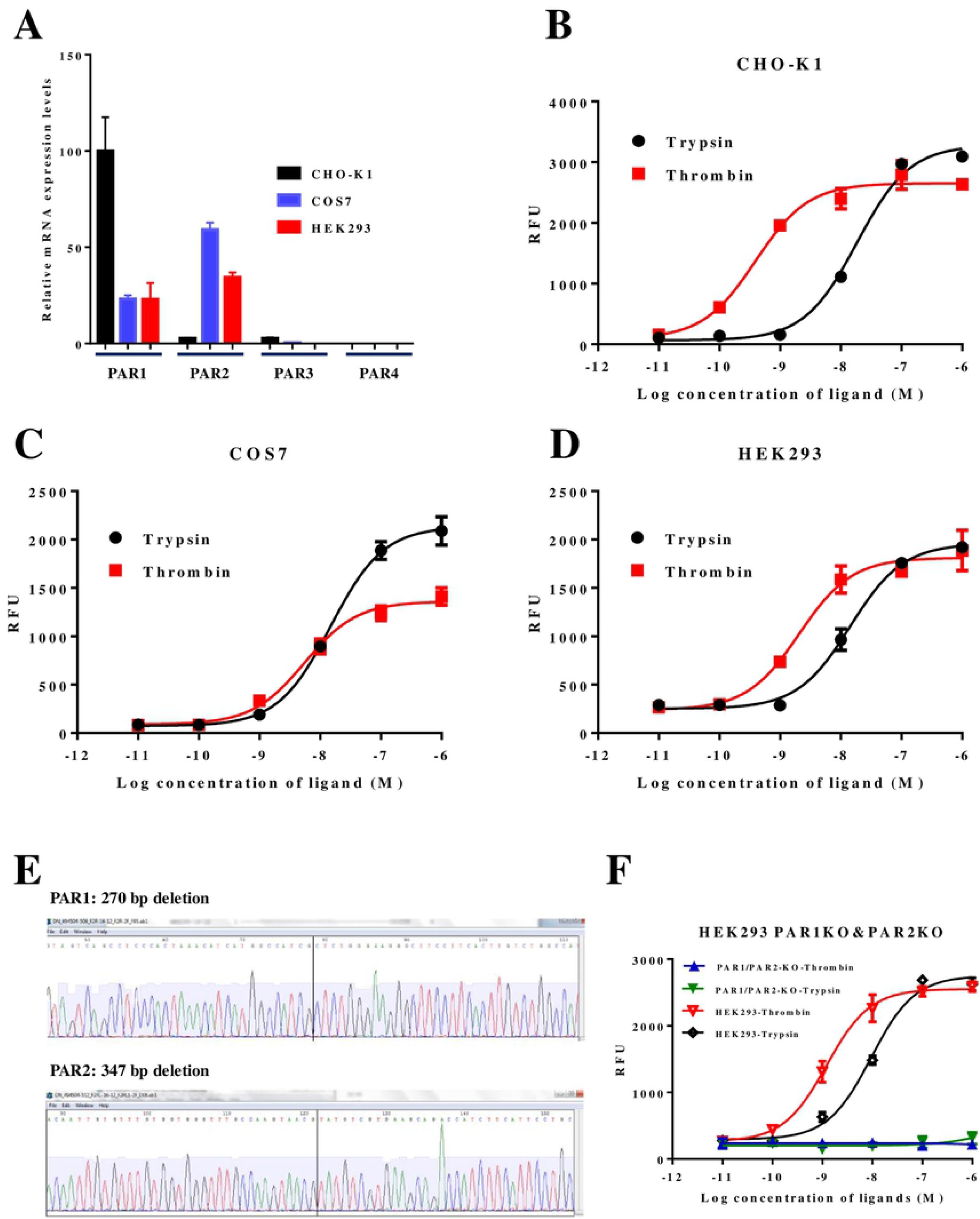
CHO-K1, COS7, and HEK293 cells express PAR1 and PAR2 receptors. A. CHO-K1, COS7, and HEK293 cells naturally express high levels of PAR1 and PAR2 mRNA but express little or no PAR3 and PAR4 mRNA. qPCR analysis was used to quantify the mRNA expression. Specific primers for each of PAR1, PAR2, PAR3, and PAR4, were used to quantify the respective mRNA expression using cDNA made from each cell line as the templates. β-actin primers were used to quantify β-actin mRNA expression as the internal controls. The relative mRNA expression of PAR1, PAR2, PAR3, and PAR4 were first normalized using β-actin expression, and then normalized using the PAR1 expression level in CHO-K1 cells, which is arbitrarily set as 100%. The relative expressions of other genes are represented as percentage of PAR1 mRNA level in CHO-K1 cells. The results shown are mean ± sd (n = 3). B, C, D. CHO-K1, COS7, and HEK293 cells naturally express PAR1 and PAR2 receptors and respond to thrombin (PAR1 ligand) and trypsin (PAR2 ligand) stimulations. FLIPR assays were used measure receptor activation as indicated by intracellular Ca^2+^ mobilization. Relative fluorescent units (RFU) are the readout for fluorescent intensities for Ca^2+^ mobilization signals. Various concentration of thrombin or trypsin were used as the ligands to activation the receptors. The assays were performed in triplicates at each data point and mean ± sd are shown. E. Sequencing analysis of the genomic DNA from *par1* & *par2* knock out HEK293 cells. The results show that a 270 bp deletion in *par1* gene and a 347 bp deletion in *par2* gene have been achieved. The deletions removed the coding regions from TM2 to TM3 for both PAR1 and PAR2 proteins. The vertical lines indicate the deletion sites. F. Characterization of *par1* & *par2* knock-out HEK293 cells. FLIPR assays were used to characterize receptor activation as indicated. Wild type HEK293 cells were used as the positive control. The assays were performed in triplicates at each data point and mean ± sd are shown.

#### Deletion of the signal peptide reduced the functional expression of PAR2, which can be rescued by a replacement signal peptide

To assess the functional role of the PAR2 signal peptide, we made a few modifications to the PAR2 N-terminus, including a N-terminal deletion to remove the signal peptide (PAR2ΔSP) and the replacement of the PAR2 signal peptide with an insulin signal peptide (PAR2-INSP), or an insulin receptor signal peptide (PAR2-IRSP) (**Fig 5A**). Pharmacological characterization of the modified receptors using FLIPR assay (**Fig 5B, C**) showed that the recombinantly expressed PAR2 responds to trypsin (EC_50_ = 1.5 nM) and PAR2 agonist peptide (PAR2-AP) (EC_50_ = 50 nM) with much higher sensitivity compared to the endogenously expressed PAR2 in HEK293 cells (EC_50_ = 10 nM for trypsin and EC_50_ = 1.5 uM for PAR2-AP). This is likely due to the over-expression of the recombinant receptor causing a super-pharmacology phenomenon (15). In this case, the EC_50_ value is a good indicator of the relative number of receptors at the cell surface. Compared to the cells expressing the wild type PAR2, cells expressing PAR2 without its signal peptide demonstrated dramatically reduced sensitivity to both trypsin and PAR2-AP (EC50 for trypsin: 50 nM; EC_50_ for PAR2-AP: 5.8 uM), suggesting the signal peptide is an important component of PAR2 functional cell surface expression. Supporting this hypothesis, replacement of the PAR2 signal peptide either with a signal peptide from insulin or from the insulin receptor fully restored the receptor ligand sensitivity (**Fig 5**).

**Fig 5.**
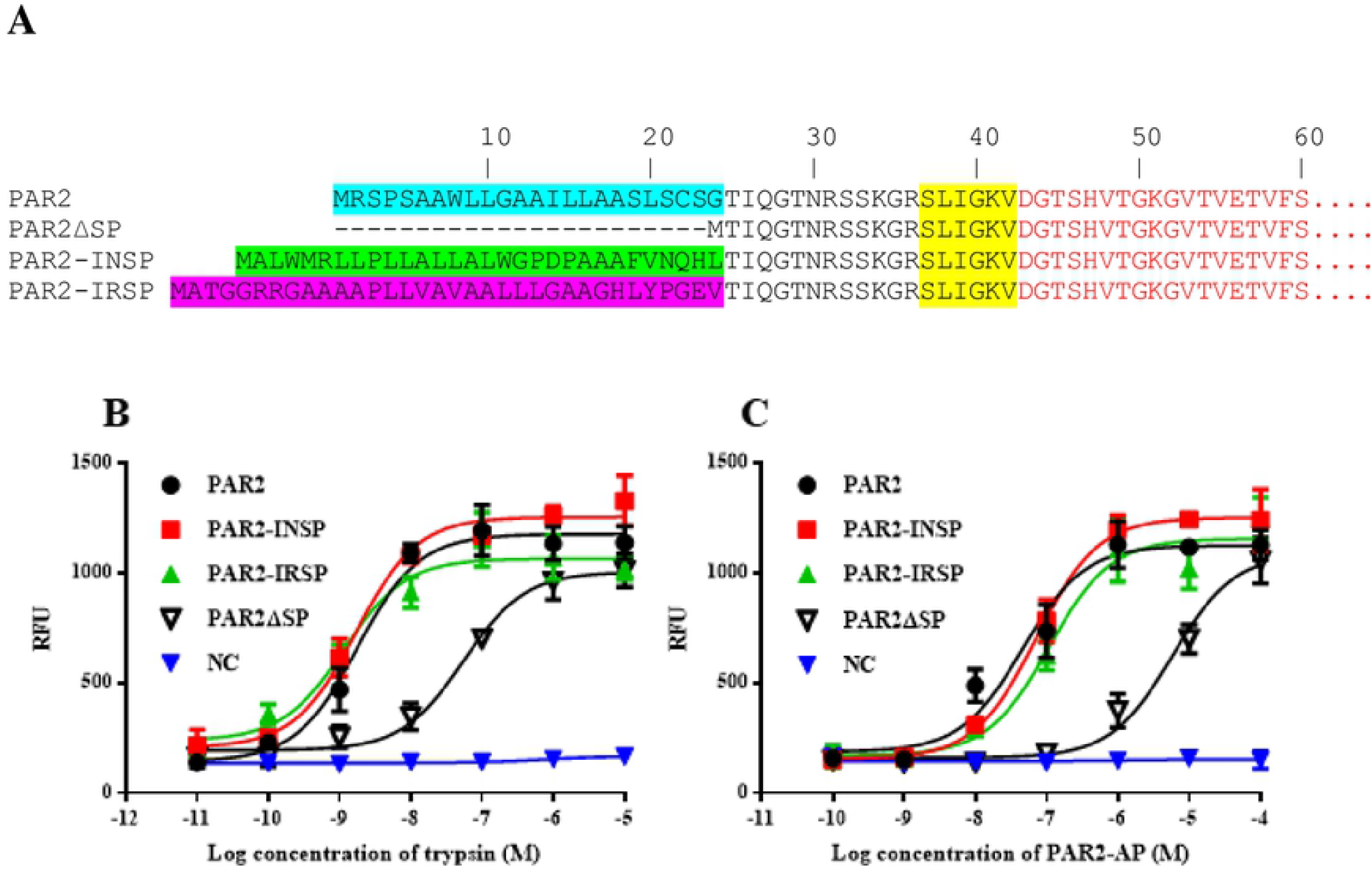
A signal peptide is important for functional expression of PAR2. A. schematic diagram showing the modifications to PAR2 receptor. The N-terminal extracellular sequences of various PAR2 mutants are shown. Human PAR2 wild type (PAR2), PAR2 with the signal peptide deleted (PAR2ΔSP), PAR2 with an insulin signal peptide (PAR2-INSP) and an insulin receptor signal peptide (PAR2-IRSP). The native signal peptide of PAR2 is shown in blue. The insulin and insulin signal peptides are shown in green and purple respectively. The tether ligand sequence of PAR2 (SLIGKV) is highlighted in yellow. B, C. Characterization of PAR2 mutants in FLIPR assay using trypsin or the synthetic PAR2 agonist peptide (PAR2-AP) as the ligands. Expression constructs for PAR2 wild type receptor and various modifications were cloned into pcDNA3.1 and transiently expressed in HEK293 cells with *par1* and *par2* knocked-out. Various concentrations of trypsin (**B**) or PAR-AP (**C**) were added to stimulate the intracellular Ca^2+^ mobilization. Relative fluorescent intensity units (RFU) are shown. The experiments were performed in triplicates at each data point and the results shown are mean ± sd. HEK293 cells with *par1* and *par2* genes knocked-out were used as the host cells for recombinant expression of various PAR2 receptors. Untransfected cells were used as the negative controls (**NC**). The experiments have been performed 3 times and very similar results have been observed. Example data is shown.

### The presence of the tethered ligand is the reason that PAR2 needs the signal peptide

#### Further deletion of the tethered ligand region rescues the functional expression of PAR2 without the signal peptide

PARs are activated by proteases, which generate new N-termini and expose the tethered peptide ligands present in the N-terminal extracellular regions of the receptors. This unique receptor activation mechanism, combined with the fact that signal peptide-less PAR2 has a poor response to ligand stimulation, led us to speculate that the necessity of the signal peptide for PAR2 may be related to the presence of tethered ligand. We made a signal peptide-less PAR2 mutant with a further deletion to the region of the tethered ligand (PAR2ΔSPΔL) (**Fig 6A**). This mutant receptor lacks the signal peptide and the tethered ligand sequence (SLIGKV) and is not activated by trypsin, however it can be fully activated by the synthetic agonist peptide PAR2-AP similarly to the wild type PAR2 receptor in the FLIPR assay (**Fig 6B**). This suggests that further deletion of the tethered ligand sequence (SLIGKV) restored functional cell surface expression of PAR2 without the signal peptide. The results also suggest that, without a signal peptide, PAR2 may be susceptible to unintended intracellular protease activation, leading to poor functional cell surface expression.

**Fig 6.**
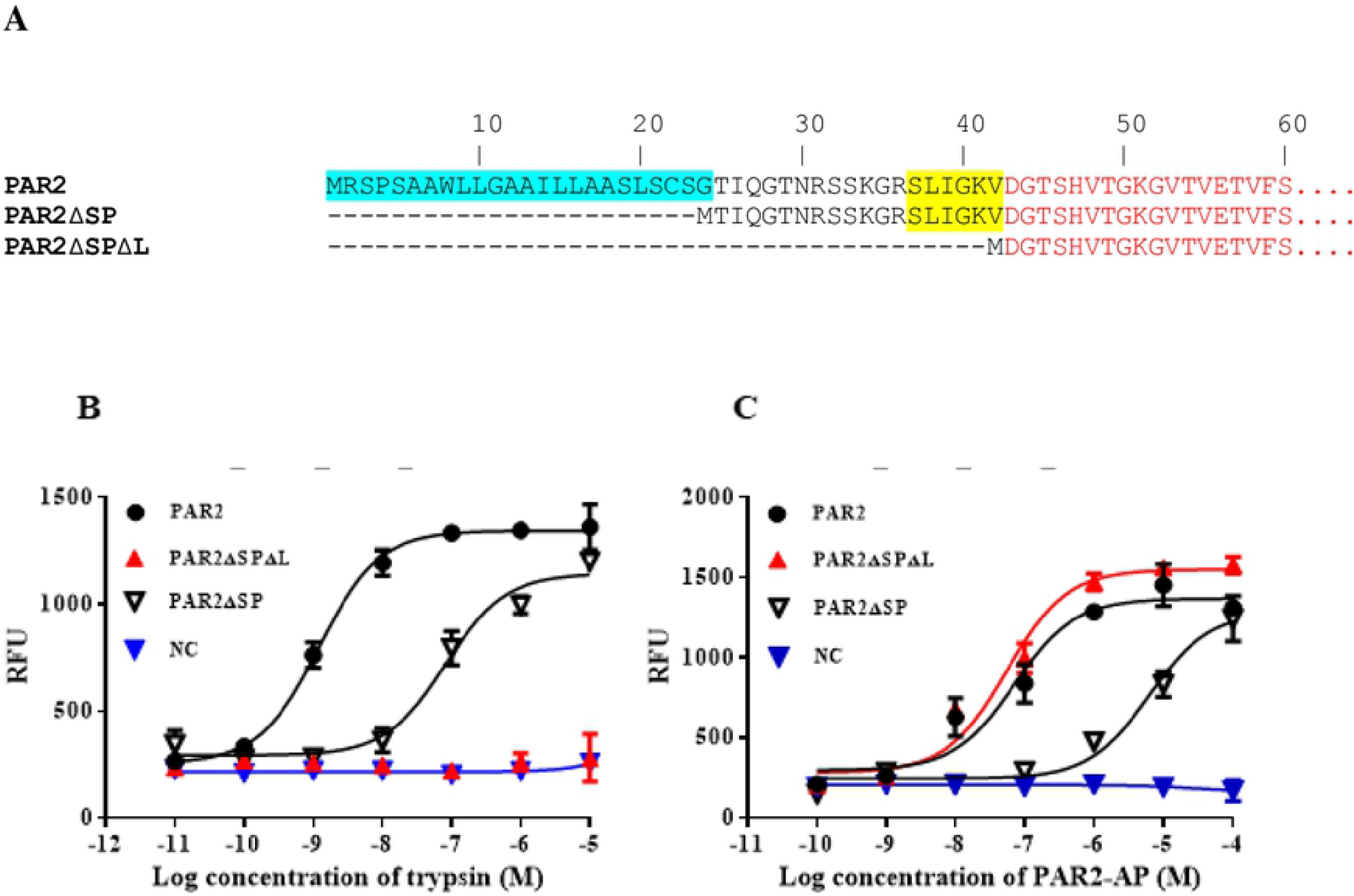
Further deletion of the tethered ligand rescues the functional expression of PAR2 without the signal peptide. A. schematic diagram showing the modifications to PAR2 receptor. The N-terminal extracellular sequences of various PAR2 mutants are shown. Human PAR2 wild type (**PAR2**), PAR2 with the signal peptide deleted (**PAR2ΔSP**), PAR2 with the signal peptide deletion and with further deletion to the tether ligand region (**PAR2ΔSPΔL**). The signal peptide of PAR2 is shown in blue. The tether ligand sequence of PAR2 (SLIGKV) is highlighted in yellow. B, C. Characterization of mutant PAR2 receptors using FLIPR assays. Various PAR2 expression constructs were transiently expressed in HEK293 with par1 and par2 knocked-out. Trypsin (**B**) or the synthetic agonist peptide PAR2 ligand (PAR2-AP) (**C**) were used as the ligand to stimulate receptor activation. HEK293 cells with *par1* and *par2* genes knocked-out were used as the host cells for recombinant expression of various PAR2 receptors. Untransfected cells were used as the negative controls (**NC**). The experiments were performed in triplicates at each data points and the results shown are mean ± sd. The experiments have been performed 3 times and very similar results have been observed. Example data is shown.

#### Mutation of Arg^36^ to Ala, which blocks the trypsin activation site, increased the functional expression of PAR2 without the signal peptide

Trypsin activates PAR2 by cleaving the protein after residue Arg^36^. This process generates a new N-terminus (with sequence SLIGKV---), which serves as a tethered ligand to activate the receptor. Mutating Arg^36^ to Ala prevents the trypsin cleavage at this position, and therefore blocks trypsin-mediated receptor activation. We made a mutation at the Arg^36^ position on PAR2 without the signal peptide (PAR2ΔSP(R36A)) and tested whether this mutation changes the level of functional receptor expression. In parallel, we also made the same mutation on the full length PAR2 receptor (PAR2(R36A)) and characterized these receptors in FLIPR assays after stimulation with trypsin or PAR2-AP. Our results demonstrated that the Arg36Ala mutation blocked, as expected, trypsin activation of PAR2 without the signal peptide (**Fig 7A**). However, when the PAR2-AP was used as the ligand, the mutant receptor (PAR2ΔSP(R36A)) demonstrated a much higher sensitivity to PAR2-AP compared to that of PAR2ΔSP (**Fig 7B**). As a control, we made the same mutation in full length PAR2 receptor (PAR2(R36A)), which responded to trypsin stimulation very poorly (**Fig 7B**), responded to PAR2-AP stimulation almost identically to the full length PAR2 receptor (**Fig 7C**). The small but detectable activation of PAR2(R36A) by trypsin (**Fig 7B**) could be due to the cleavage of PAR2 by trypsin at Arg^31^, or Lys^34^ positions, resulting in tethered ligands with poor activity for receptor activation.

**Fig 7.**
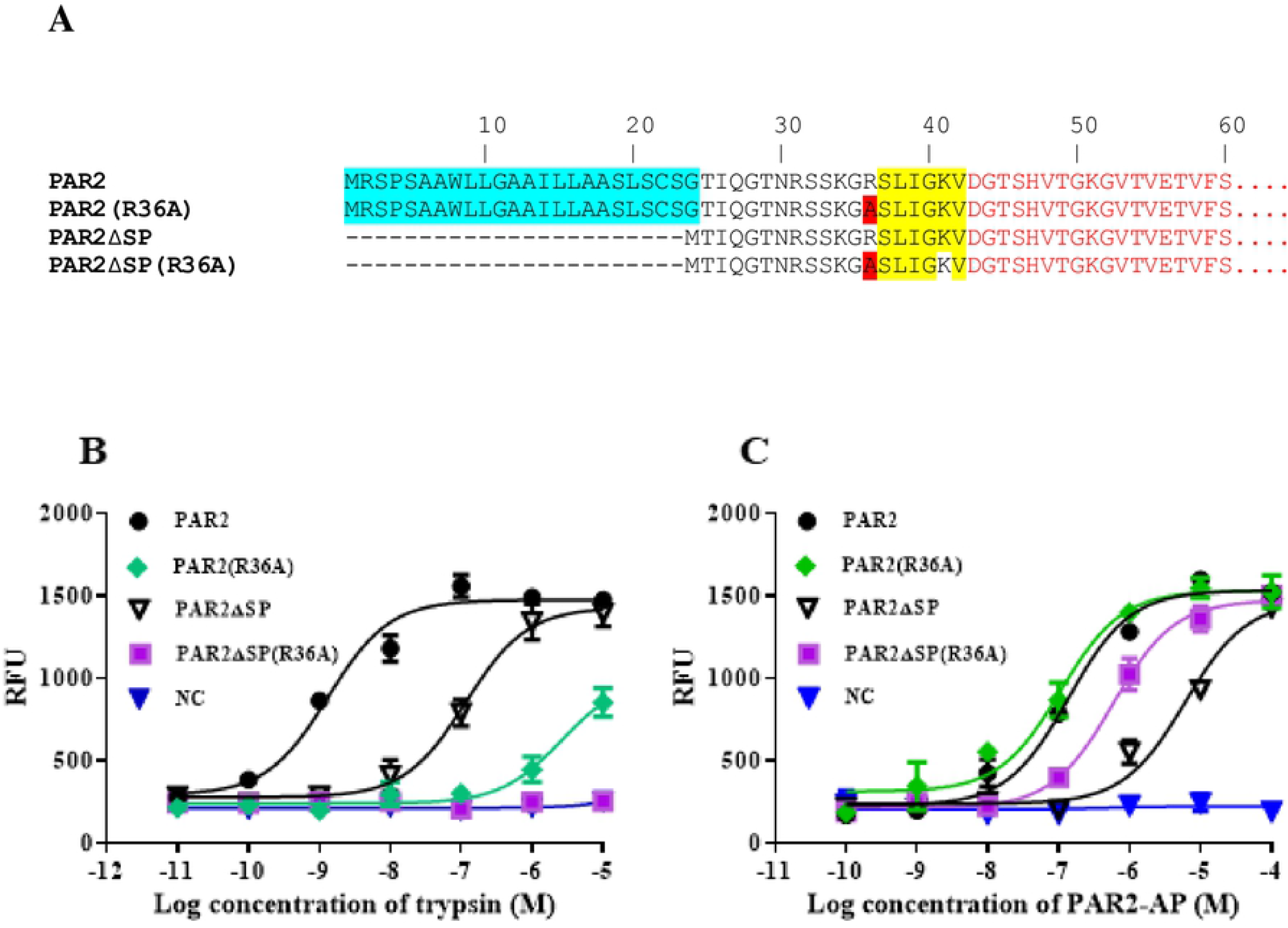
Arg^36^ to Ala mutation helps the functional expression of PAR2 without a signal peptide. A. schematic diagram showing the modifications/mutations to PAR2 receptor. The N-terminal extracellular sequences of various PAR2 mutants are shown. PAR2 wild type (**PAR2**), PAR2 with a Arg^36^ mutation to Ala (**PAR2(R36A)**), PAR2 with the signal peptide deleted (**PAR2ΔSP**), PAR2 with the signal peptide deletion and with an Arg36Ala mutation (**PAR2ΔSP(R36A)**) were used for characterizations. The signal peptide of PAR2 is shown in blue. The tether ligand sequence of PAR2 (SLIGKV) is highlighted in yellow. The mutation Ala residue from Arg^36^, which is involved in trypsin cleavage/activation of PAR2, is highlighted in red. The mutant receptors were characterized in FLIPR assays using either trypsin (**B**) or PAR2-AP (**C**) as ligands. HEK293 cells with *par1* and *par2* genes knocked-out were used as the host cells for recombinant expression. Untransfected cells were used as the negative controls (**NC**). The experiments were performed in triplicates at each data points and the results shown are mean ± sd. The experiments have been performed 3 times and very similar results have been observed. Example data is shown.

#### Protease inhibitor treatment increased functional expression of PAR2 without the signal peptide

We hypothesized that serine protease inhibitors can help the functional expression of PAR2 without a signal peptide by blocking premature intracellular protease-mediated activation. A protease cocktail including AEBSF, Leupeptin, and aprotinin was used to inhibit ER and Golgi proteases (16, 17). Cells expressing the wild type PAR2 and various mutant forms of PAR2 were treated with the protease inhibitor cocktail and then tested for their responses to PAR2-AP stimulations. Trypsin was not used in this assay because it is inhibited by the protease inhibitor cocktails. Our results showed that while protease inhibitors did not affect the EC_50_ values of PAR2-AP stimulated responses for PAR2 wild type, PAR2(R36A), PAR2ΔSP(R36A), and PAR2ΔSPΔL, the protease cocktail clearly increased functional expression of PAR2ΔSP by decreasing the EC_50_ value (from 5.8 uM to 0.7 uM) (**Fig 8**).

**Fig 8.**
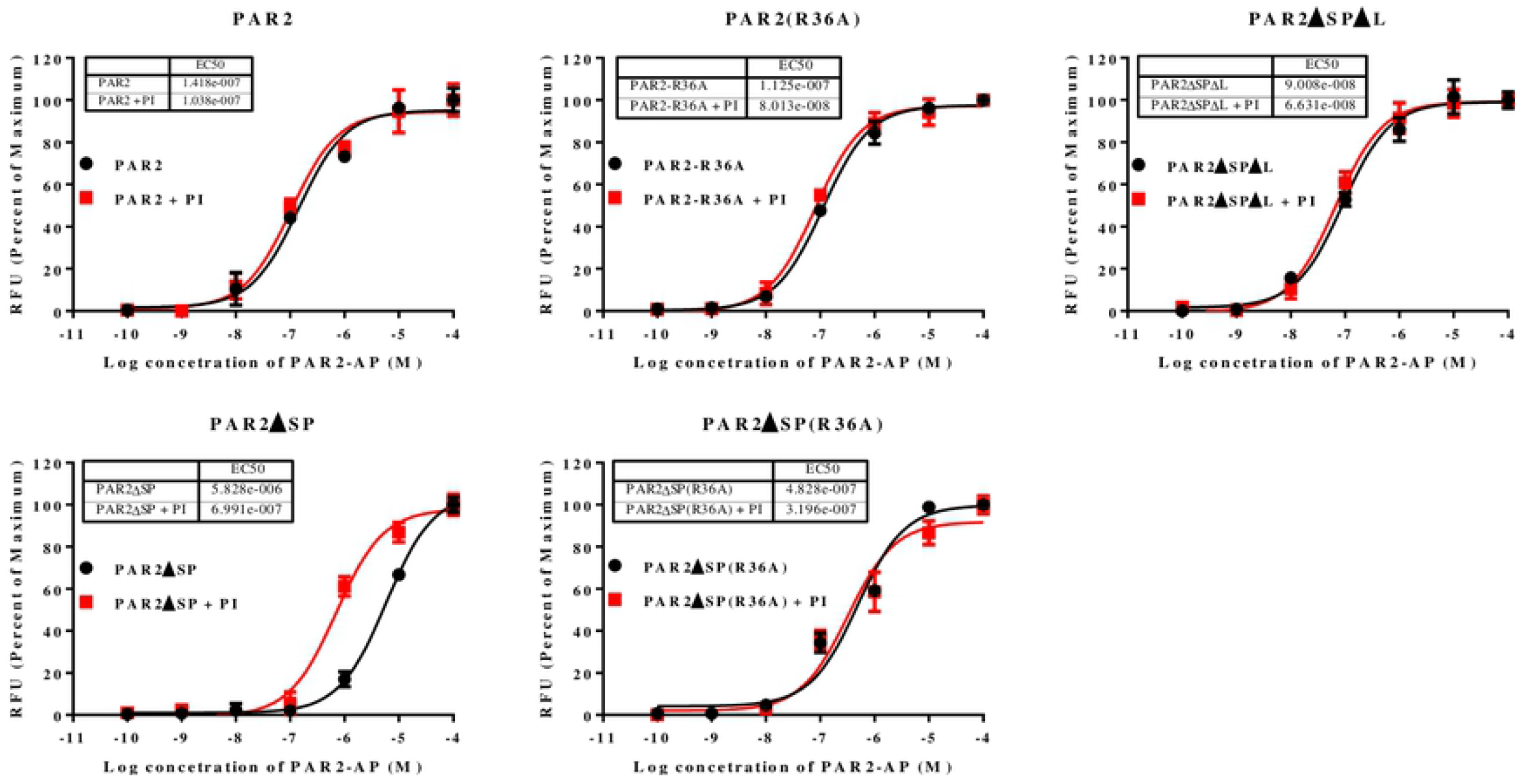
Serine protease inhibitor cocktail increases the functional expression of PAR2 without the signal peptide. HEK293 cells with *par1* and *par2* knocked out were used for the transient expression of various PAR2 proteins. Treatment with proteases inhibitor cocktail (**PI**) lowered the Emax values for all receptors with similar degrees. Protease treated samples showed about 80% response in Emax values compared with those of untreated cells. For good comparison of the EC50 values between samples treated and untreated with the protease inhibitor cocktails, the results were normalized using their Emax values and the data were expressed as the percentages of the Emax. The experiments were performed in triplicates at each data points and the results shown are mean ± sd. The experiments have been performed 3 times and very similar results have been observed. Example data is shown.

#### Arg36Ala mutation and the protease inhibitor treatment increase the cell surface expression of signal peptide-less PAR2

To confirm whether the reduced responses of signal peptide-less PAR2 to the ligand stimulation is due to a lack of total receptor protein expression, and/or a lack of cell surface expression, we used a monoclonal antibody against the amino acid residues 37-62 of PAR2 in ELISA assays to measure the total and cell surface expression of the various forms of PAR2 and effect of protease inhibitor treatment. We observed that PAR2 wild type and PAR2(R36A) mutants have the highest total and cell surface protein expression as measured by ELISA. PAR2ΔSPΔL has slightly lower expression compared to that of the PAR2 wild type in both total and cell surface expression. As this variant of PAR2 is missing amino acid residues 1-42, its reduced detection of proteins expression could be due to the poor antibody recognition. PAR2ΔSP(R36A) has further lower total and cell surface expression and PAR2ΔSP has the lowest total and particularly cell surface expression levels (**Fig 9**). The data showed that the great majority of PAR2ΔSP protein are located intracellularly and only a small portion of it are present on the cell surface. For PAR2 wild type, PAR2(R36A), and PAR2ΔSPΔL, over 90% of the proteins are presented on the cell surface. Corroborating the functional assays, protease inhibitor treatments increased the total and especially the cell surface expression levels for PAR2ΔSP while having little or no effects on the protein expression of other forms of PAR2 proteins (**Fig 9**). Stimulation of receptors using PAR2 peptide ligand (PAR2-AP) decreased the cell surface and the total protein expression levels for all variants of PAR2 except PAR2ΔSP. This is likely due to that the majority of PAR2ΔSP being intracellular, and the ligand stimulation of the cell surface receptor, causing the subsequent internalization and degradation of the stimulated receptors applies less to PAR2ΔSP.

**Fig 9.**
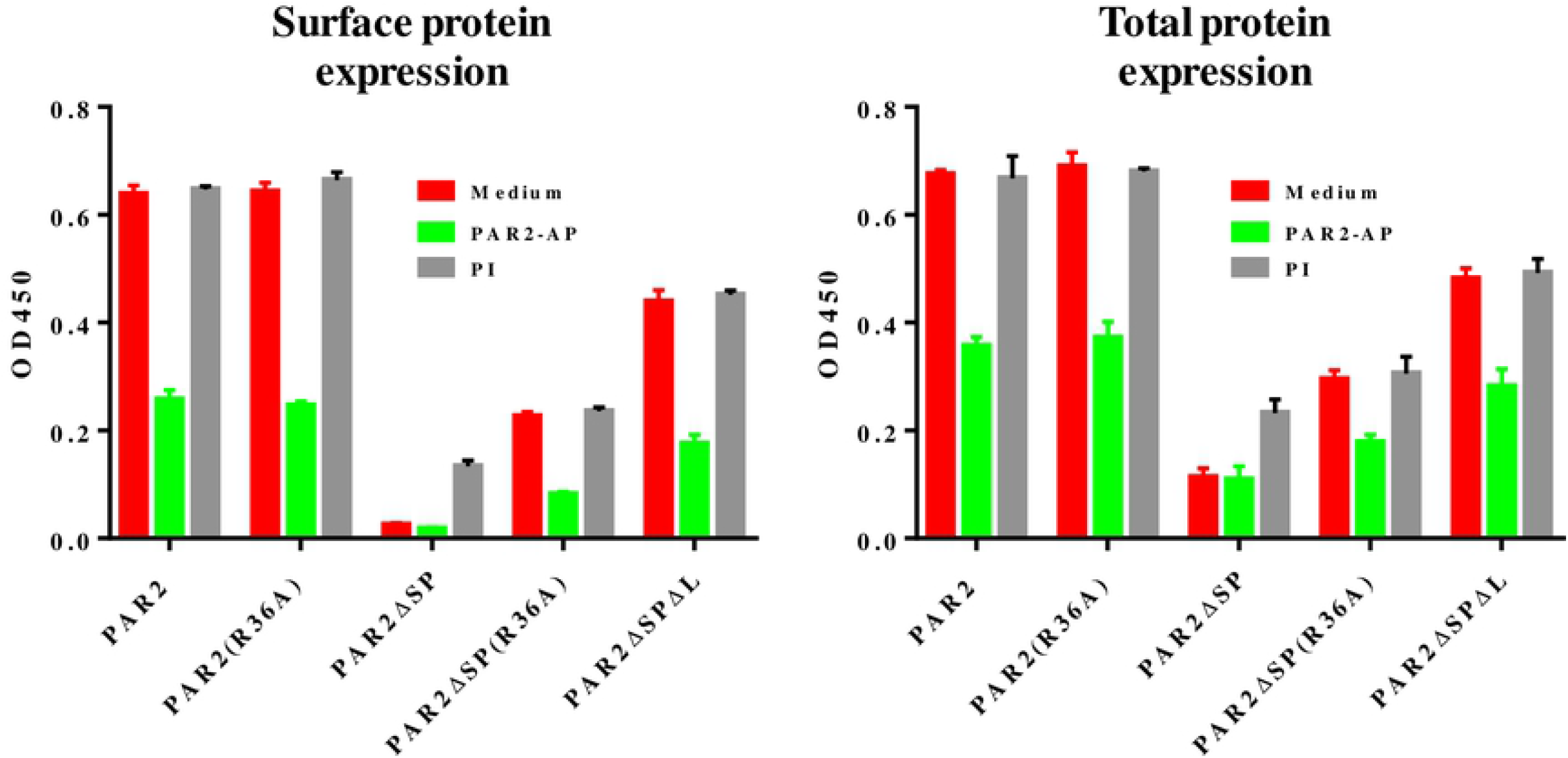
Cell surface and total protein expression of PAR2 wild type and mutants. HEK293 cells with *par1* and *par2* knocked-out were used for the transient expression of various PAR2 proteins. PAR2 peptide ligand, PAR2-AP and protease inhibitor cocktails (**PI**) were used for treatments. Medium was used as the control treatment. ELISA with or without cell penetrating reagent was used to measure the total and cell surface and protein expression. The experiments were performed in triplicates at each data points and the results shown are mean ± sd. The experiments have been performed 3 times and very similar results have been observed. Example data is shown.

In parallel, to further facilitate the measurements of the protein expression and visualization of protein cellular localizations, we constructed various PAR2 expression vectors by fusing a GFP tag to the C-termini of the PAR2 wild type protein and the various PAR2 mutants (**Fig 10A**) and expressed them in the *par1* and *par2* null HEK293 cell line. We measured the total expression levels of PAR2 and the mutant proteins by measuring GFP fluorescence intensity of the various GFP fusion proteins. In general, the results are very similar to that shown by ELISA using the anti-PAR2 antibody except for PAR2ΔSPΔL, which showed lower levels compared to that of PAR2 wild type in the ELISA assays, but showed similar expression levels to that of PAR2 wild type in GFP intensities (**Fig 10B**). This result supports our earlier speculation that the reduced detection of PAR2ΔSPΔL expression is likely due to the poor recognition of PAR2ΔSPΔL (which misses the amino acid residues 37-42 of the recognition site) by the antibody. To investigate the cellular localizations of PAR2 protein and its variants, we used confocal microscopy to analyze the cells that express various PAR2 proteins at various conditions including the treatments with PAR2 ligand or protease inhibitors. PAR2 wild type, PAR2(R36A), and PAR2ΔSPΔL proteins are mainly localized on the plasma membranes (**Fig 10C**). PAR2ΔSP can only be found intracellularly and little or none is on the plasma membranes, which is very similar to PAR2 wild type receptor stimulated by the peptide ligand (PAR2+PAR2-AP, **Fig 10C**). For PAR2ΔSP(R36A), a good portion of protein is expressed on the plasma membrane and a large amount of protein can also be found intracellularly. Interestingly, protease inhibitor treatment enabled the plasma membrane expression of PAR2ΔSP (PAR2ΔSP+PI, **Fig 10C**). Overall, the observed GFP-tagged protein cellular distribution is in agreement with the ELISA data (**Fig 9**). Interestingly, the cells expressing PAR2ΔSP(R36A) and protease inhibitor-treated cells expressing PAR2ΔSP appeared to belong two subcategories. One population of cells have good PAR2 plasma membrane localization, mimicking the wild type PAR2, and another population of cells only have intracellular PAR2 which is similar to that of PAR2ΔSP without the protease inhibitor treatment. Arg36Ala mutation and the protease inhibitor cocktails (used in the assays) mainly block PAR2 cleavage/activation by the serine proteases. However, cells may express other proteases that can cleave and activate PAR2ΔSP intracellularly but would not be blocked by the mutation or the protease inhibitor treatment. It is possible that cells express different proteases under different conditions such as different cell cycle stages (18–25).

**Fig 10.**
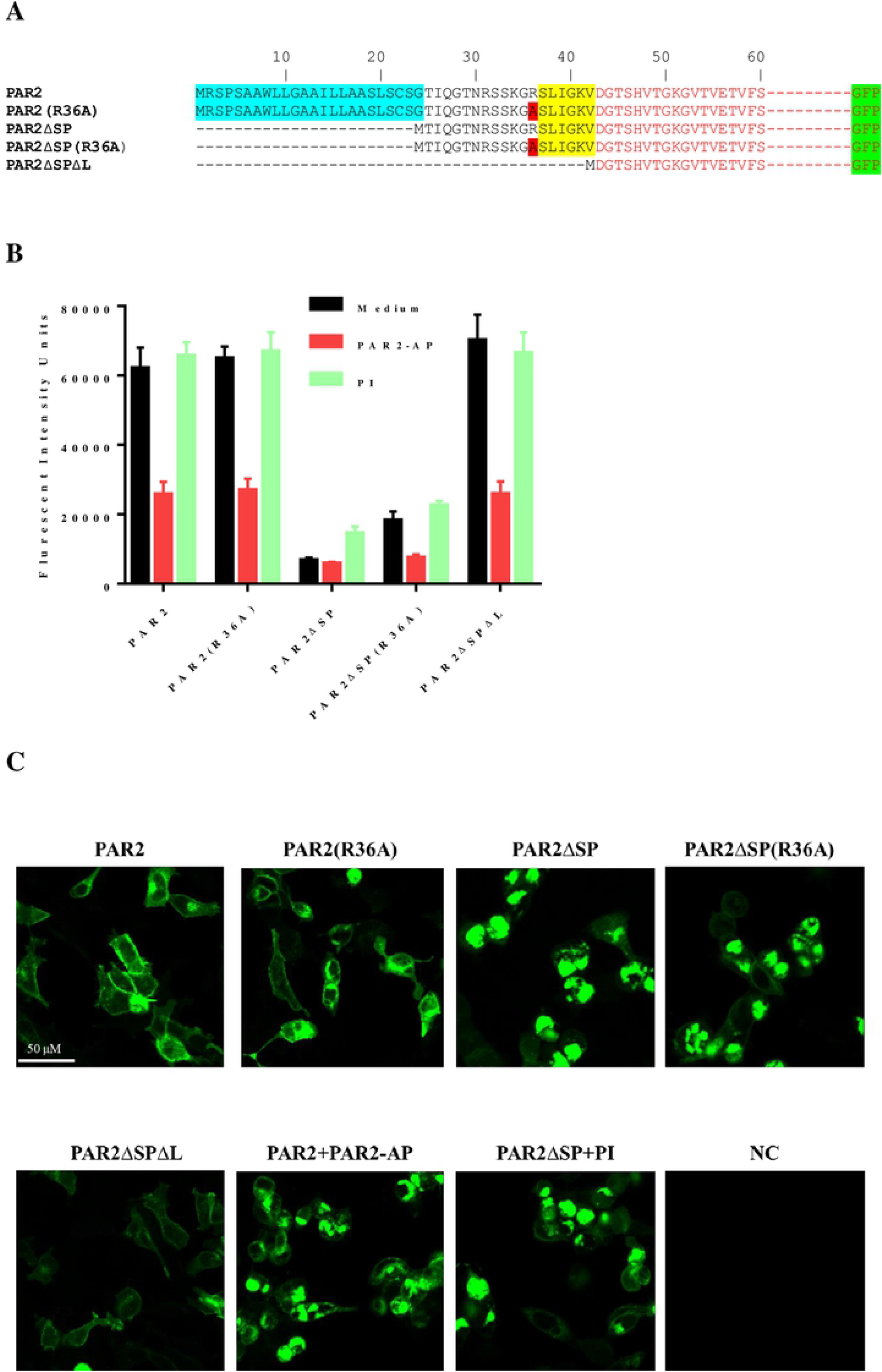
Arg36Ala mutation and protease inhibitors increase the cell surface expression of PAR2-GFP without a signal peptide. **A. Schematic presentation of various PAR2-GFP fusion protein expression constructs. B. The expression levels of various PAR2-GFP proteins with treatments of PAR2-AP, or protease inhibitors**. Various PAR2-GFP expression constructs were transiently expressed in HEK293 cells with *par1* & *par2* knocked-out. The transfected cells were treated either with medium (**medium**), peptide agonist (**PAR2-AP**), or a protease inhibitor cocktails (**PI**), and the fluorescent intensities of the cells expressing the PAR2-GFP fusion proteins were measured. Assays were performed in quadruplicates at each data point. **C. Fluorescent images from confocal microscope showing the cellular distributions of various PAR2-GFP fusion proteins under the treatments of PAR2-AP or protease inhibitors**. Untransfected cells were used as the negative control (**NC**). The fluorescent intensities shown in the images are automatically adjusted for better viewing of the protein cellular distributions. The experiments have been performed 3 times and very similar results have been observed. Example data is shown.

GPCRs are synthesized in the endoplasmic reticulum (ER) and transported to the Golgi apparatus and then to the plasma membrane. There are many proteases present in the endoplasmic reticulum and Golgi apparatus (16, 26–29) which may cleave the protease-sensitive PAR2 activation site at Arg^36^ position during the protein synthesis and transportation process. This would cause unintended or premature receptor activation, which would subsequently lead to receptor internalization and degradation. The signal peptide of PAR2 is important for its functional expression. However, the removal of the tethered ligand or the blockage of the receptor activation by proteases dismissed the necessity of the signal peptide, suggesting that the signal peptide may help prevent this unintended cleavage of PAR2 at the activation site during the protein synthesis and/or transportation process. For cell surface proteins using signal peptides, their translocation to ER and eventually the plasma membrane is mediated by ER translocons (30, 31), which play roles in protein compartmentalization (32–36) and segregation (31, 37, 38). Although the mechanism remains unclear, we speculate that ER translocons may play the role in protecting PAR2 from protease cleavage (**Fig 11**).

**Fig 11.**
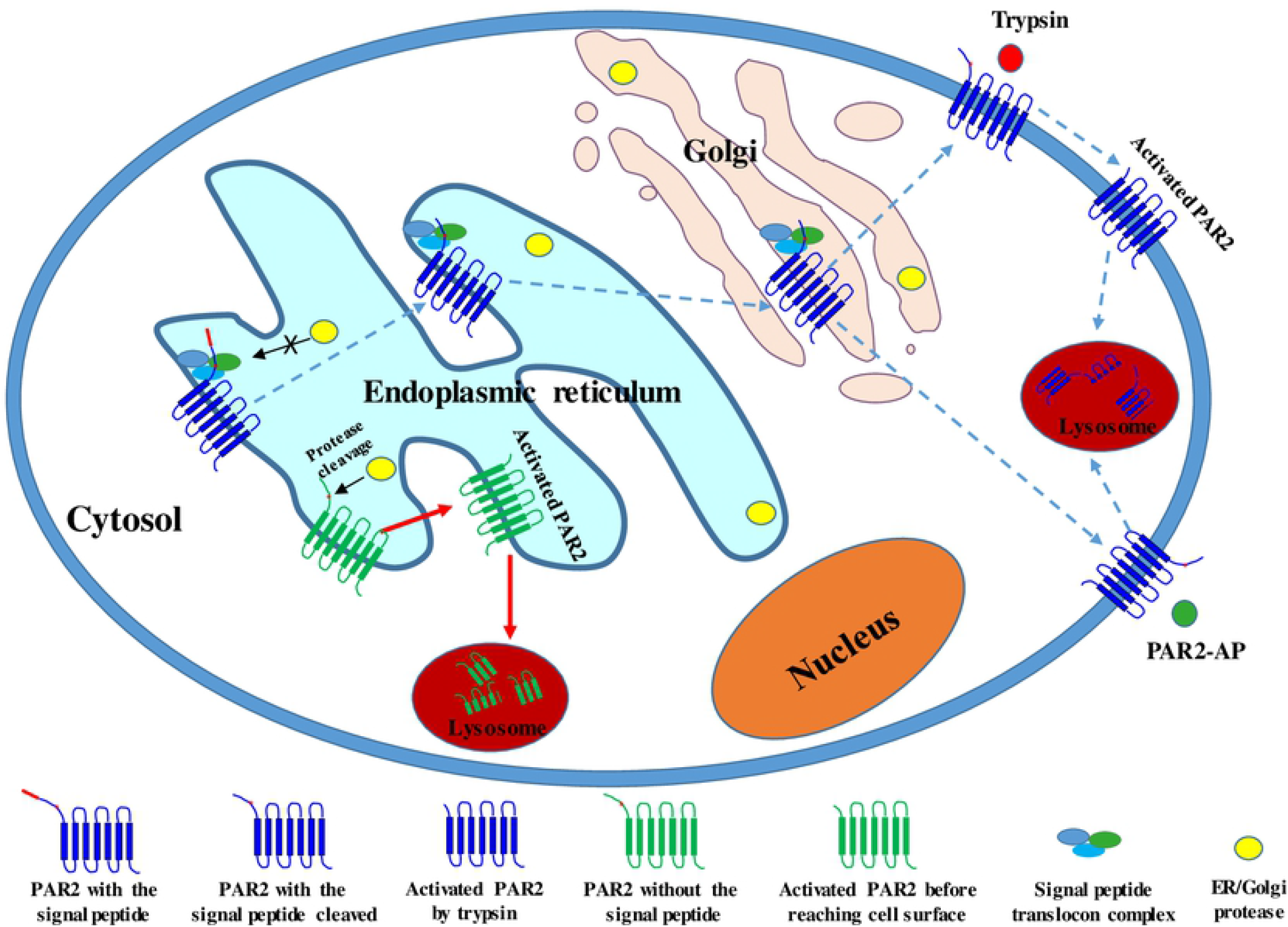
Schematic diagram showing the proposed role of PAR2 signal peptide in protecting PAR2 from protease cleavage before reaching the plasma membrane. Without the signal peptide, the protease activation site of PAR2 is susceptible to protease cleavage in ER and Golgi, leading to PAR2 activation before reaching the cell surface and subsequent translocation to lysosome for degradation. With the signal peptide, PAR2 is bound by the signal peptide related translocon complex and segregated/protected from the cleavage by ER/Golgi proteases, allowing the receptor to reach the plasma membrane for sensing the extracellular trypsin activation. The signal peptide of PAR2 at the N-terminus is shown in red.

The classical signal peptide has been known to help secreted proteins and cell surface proteins to cross or become embedded in the cell membranes. In this report, through studying the signal peptide of PAR2, we observed a function of the signal peptide to serve as a protector of PAR2 from intracellular protease activation. Based on the high sequence similarities among the PARs and their similar activation mechanisms, the signal peptides for PAR1, PAR3, and PAR4 may also serve the same function as that of PAR2. Cleavage of PARs by intracellular proteases leads to the unintended activation of the receptors and the loss of function to sense the extracellular signals. Therefore, with the protease-protection function, the signal peptides are critical for the functions of the PAR family of receptors.

## Summary

To summarize, in this report we observed that deletion of the signal peptide of PAR2 decreased PAR2 cell surface expression with the most receptors accumulating intracellularly. However, further deletion of the tether ligand of PAR2, which disables the activation of PAR2 by trypsin, restored the receptor cell surface expression, suggesting that the necessity of the signal peptide for PAR2 is related to the presence of the tether ligand sequence and the protease activation mechanism. We hypothesize that the signal peptide of PAR2 protects PAR2 from intracellular protease cleavage and activation. Without the signal peptide, PAR2 can be cleaved and activated by intracellular proteases in the endoplasmic reticulum or Golgi apparatus, leading to the unintended, premature receptor activation and resulting in intracellular accumulation. Supporting this hypothesis, an Arg36Ala mutation at the trypsin activation site as well as protease inhibitor treatments both increased the cell surface expression of the signal peptide-less PAR2 and functional responses to ligand stimulation. Our results extend the knowledge of PAR2 expression/function and reveal a role of the signal peptide in protecting cells surface protein, and perhaps the secreted proteins as well, from intracellular protease cleavages.

